# Sexual conflict, directional sexual selection and phenotypic plasticity jointly drive the evolution of extreme phenotypic variation

**DOI:** 10.64898/2026.08.18.745420

**Authors:** Claudia Pruvôt, Arnaud Badiane, Ingrid Dourlens, Mariama Dramé, Juliette Mendes, Mirjam Urb, Romain Vedie, Severine Viala, Cristinia Vieira, Patricia Gibert, Abderrahman Khila

**Affiliations:** Institut de Génomique Fonctionnelle de Lyon, CNRS UMR5242, École Normale Supérieure de Lyon, Université Claude Bernard Lyon1, Lyon France; Laboratoire de Biométrie et Biologie Evolutive, Université Claude Bernard Lyon 1, CNRS, Villeurbanne, France

## Abstract

How broad phenotypic variation is maintained in natural populations in the face of selection is a central question in evolutionary biology. We address this question in the water strider *Microvelia longipes,* where males exhibit striking variation in rear leg length used in male-male contests for dominance. Using reaction norm experiments on inbred lines, we demonstrate that phenotypic plasticity contributes to expanding phenotypic variation, but requires high genetic variation to generate the broad range of trait expression observed in natural populations. Experimental evolution favouring trait exaggeration revealed that directional sexual selection not only fails to erode variation of male rear leg length, but rather amplifies it beyond the natural distribution. Additionally, male-limited selection in favour of dominance generated substantial fecundity costs in females, underscoring the role of sexual conflict driven by females in constraining exaggerated secondary sexual traits in males. Our findings show that sexually antagonistic selection and directional sexual selection jointly generate high genetic variation, which phenotypic plasticity inflates into broad phenotypic distribution of male weapon size. This provides an empirical explanation for the high variability of male exaggerated weapons in nature.

## Introduction

Exaggerated secondary sexual traits are widespread (Miller et al., 2026) and often display extraordinary levels of phenotypic variation within populations (Kotiaho, 2001; Pomiankowski and Moller, 1995). In addition to being highly variable, these traits are typically condition-dependent, such that their expression is strongly influenced by environmental factors experienced during development (Andersson and Iwasa, 1996). Consequently, the remarkable variation observed in sexually selected traits is thought to arise from the combined effects of genetic variation and phenotypic plasticity.

The persistence of such extensive variation nevertheless represents a long-standing question in evolutionary biology. Exaggerated secondary sexual traits are generally subject to strong directional sexual selection, which is predicted to drive the underlying alleles towards fixation thus reducing additive genetic variance (Falconer and Mackay; Kaufmann et al., 2023; Lande, 1980; Lynch and Walsh, 1998). Yet substantial phenotypic variation is maintained within natural populations, suggesting that additional evolutionary processes counteract the erosion of genetic variation expected under persistent directional selection.

Several works, both theoretical and empirical, described mechanisms such as mutation-selection balance or sexually antagonistic selection that counteract the effect of directional selection to maintain genetic variation (Kaufmann et al., 2023; Kirkpatrick and Ryan, 1991; Pomiankowski and Moller, 1995; Rowe and Houle, 1996). The genic capture model, for example, proposes that sexually selected traits reflect an individual’s overall condition, which is itself determined by a large number of loci affecting resource acquisition, metabolism and allocation, and, therefore, they are expected to indirectly capture genome-wide genetic variation (Rowe and Houle, 1996). Consistent with this prediction, experimental evolution in bulb mites showed that selection favouring exaggerated male traits captures genetic variation across the genome while simultaneously reducing overall genomic diversity (Parrett et al., 2022). Other studies however came to different conclusions. Experimental tests of the role of condition-dependence in maintaining genetic variance of multiple male sexually selected traits in the fly *Drosophila bunnanda* found condition-dependent expression to be insufficient to maintain genetic variance available to sexual selection in these traits (Van Homrigh et al., 2007). Sexually antagonistic selection, on the other hand, can preserve genetic diversity when alleles increasing male reproductive success reduce female fitness, preventing their fixation within populations (Chippindale et al., 2001; Connallon and Clark, 2014; Kaufmann et al., 2023). Likewise, genotype-by-environment interactions and phenotypic plasticity may amplify standing genetic variation by generating a broad spectrum of trait expression across environmental conditions (Kokko and Heubel, 2008). Despite considerable progress, studies that address the interaction between these mechanisms remain difficult to achieve. Understanding how various facets of selection jointly maintain high genetic variation and how phenotypic plasticity contributes to amplifying this variation is therefore essential for understanding how weapons of competition come to be so variable among males in the face of directional sexual selection operating on these traits.

Here, we investigate the various mechanisms contributing to the evolution of extreme variation of an exaggerated sexually selected trait in the males of the water strider *Microvelia longipes* (Toubiana and Khila, 2019). Males of *M. longipes* display striking variation in the length of their rear legs, which function as weapons during male–male competition (Toubiana and Khila, 2019). Previous studies have shown that males with elongated legs are more successful in dominating egg-laying sites, securing copulations with gravid females, and siring the majority of offspring (Toubiana and Khila, 2019). While these findings implicate directional sexual selection in favouring the long-legged males, the persistence of small and intermediate males accounting for the striking variation in leg length remains unexplained. We show that phenotypic plasticity alone fails to explain the broad variation observed in male rear leg length. Experimental evolution favouring exaggeration revealed that directional sexual selection not only fails to erode variation of male weapon, but rather is necessary to maintain it. Furthermore, directional sexual selection favouring male weapon also imposes fecundity costs on females, highlighting the joint role of sexually antagonistic selection and directional sexual selection in generating variation in this trait.

## Results

### Alone, phenotypic plasticity fails to explain the broad variation in male leg length

The extent to which condition dependence expands standing genetic variation into extreme leg length variation in the males of *Microvelia longipes* is unclear. We therefore conducted a series of experiments aimed at isolating the effect of nutrition-driven plasticity from that of genetic variation on male leg length variation. Distinct natural populations of *Microvelia longipes,* from French Guiana (FG) and Brazil, showed high and consistent male leg length variation with a coefficient of variance of 25 % and 29% respectively, and a mean male leg length of 6439µm and 6542µm (**Figure 1**). The coefficient of variance of body size in the FG and Brazil populations was 7.8% and 8.8% respectively. The French Guiana population was maintained in the laboratory for over 10 years with no external gene flow, thus reducing genetic variation. Under these laboratory conditions, sexual selection favouring long legged males was reduced or absent (see material and methods for detail) and the coefficient of variation dropped from 25% to 15% and 17% in rich and poor diet respectively (**Supplementary table S1**). This observation prompted us to further test the combined effect of high genetic variation and phenotypic plasticity in generating extreme male leg length variation in *Microvelia longipes*.

**Figure 1.**
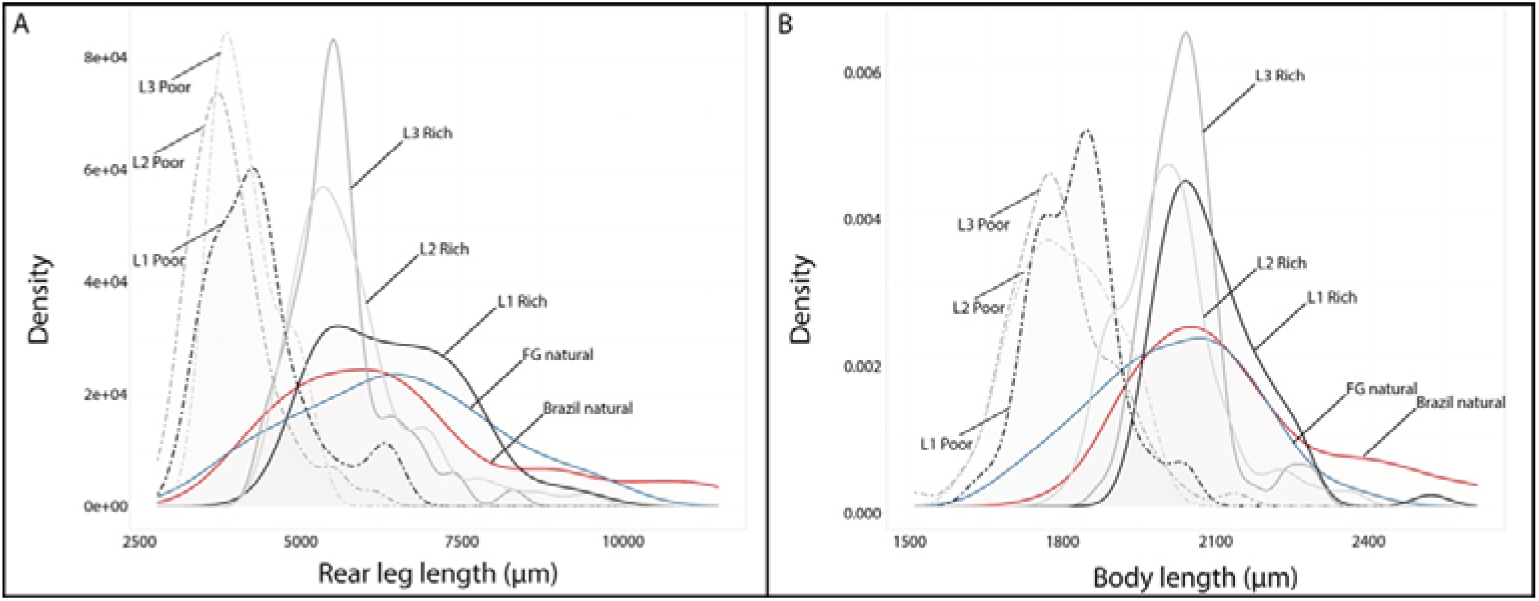
Effect of plasticity and genetic variation on the variation rear leg size (**A**) and body size (**B**) in the males of *Microvelia longipes*. Red represents a wild population collected in Brazil in 2015 and blue a wild population collected in French Guiana in 2013. Light grey, dark grey and black represents three inbred lines derived from the French Guiana population. Continuous lines represent rich diet and dashed lines poor diet treatments.

Next, we examined leg length variation in three inbred lines derived from the French Guiana population and generated through 20 generations of brother-sister crosses of randomly chosen males at each generation (Toubiana and Khila, 2019). The coefficients of variance of male leg length and body size of these lines dropped dramatically, in both poor and rich diet, and in some cases by over half in comparison with the initial natural population (**Figure 1; Table S1**). Therefore, while phenotypic plasticity contributes to expanding male leg length variation to a certain extent, it is not sufficient alone and requires the presence of high standing genetic variation to generate extreme variation in this trait.

### Selection favouring dominant males is necessary to maintain variation of male weapon

Next, we sought to identify the mechanisms responsible for the persistence of both small and large males in the population. Long-legged males dominate egg-laying sites and sire most of the eggs laid on these sites (Toubiana and Khila, 2019). If the accumulation of alleles associated with long legs is driven solely by selection favouring dominant males, we should expect an increase in mean male leg length and a decrease in the variation of this trait in populations subjected to directional sexual selection in favour of exaggeration. Conversely, we should expect mean leg length to decrease and the variance to remain high in populations under relaxed selection. To test these hypotheses, we conducted a large-scale experimental evolution study designed to reduce the effects of phenotypic plasticity while imposing strong directional selection favouring long-legged males (see methods for detail). As controls, we created lines where male-male competition is relaxed to absent (see methods).

Rear leg and body length variation in both sexes was similar between the wild population and its F1 progeny that we used as the base population, for both males and females, indicating that there was no loss of phenotypic variation during the expansion of the population (**Figure 2**). At the start of the experiment, males from both control and selection lines showed no difference in leg or body length (leg length p = 0.732, body length p = 0.931) (**Figure 2A-B**). This also indicates that there was no bias when lines and replicates were created. At G1, males from the control line (raised on richer diet) had rear legs 8.6% longer (p < 0.0001), while males from the selection line (raised on less rich diet) had overall smaller legs and bodies (p < 0.0001) when compared to the base population (**Figure 2A-B**). This is consistent with the differences in nutritional treatment applied to the control and selection lines (see methods for detail). Between generations 5 and 10, mean male rear leg length in the selection lines caught up and exceeded that of control lines even though control lines were raised on richer diet (**Figure 2A**). At generation 10, males from the selection line (fed on standard less rich diet) had rear legs 17.2% longer (p < 0.0001) than males from the control line (fed on richer diet). This difference between the experimental lines persisted at generation 15 (**Figure 2A**). Although mean length of male rear legs in the selection lines increased over generations, it did not exceed that of the base population at generation 15 (p = 0.0514), but this effect is under-estimated due to differences in nutritional treatment. We conclude that the divergence between the lines is primarily due to the decrease of the mean in the control line, in which males from generation 15 had rear legs that were 11.8% shorter than the base population (p < 0.0001) (**Figure 2**). Surprisingly, the effect of experimental evolution on male rear leg length variation was contrary to our expectations. At generation 15, variation in male rear leg length had increased by 16% in the selection line (p = 0.0164) and had decreased by 47% in the control line (p < 0.0001) compared to the base population (**Figure 2**). In both lines, the mean of male body length showed the same pattern as rear leg length, consistent with previous work establishing that these two traits covary (Toubiana et al., 2021). However, the effect size on the body is much smaller than for the length of the rear legs. Specifically, the difference in body length between males from the selection and the control lines was about 4% (p < 0.0001) at generation 15 (**Figure 2B**). This indicates that body length responded to selection favouring dominant males with a lower rate compared to rear leg length. Contrary to rear leg length, body length variation did not differ between generations in either line (selection line p = 0.3962, control line p = 0.5784), although there was a 19% difference in body length (p < 0.0001) between the selection and the control lines at generation 15 (**Figure 2B**).

**Figure 2.**
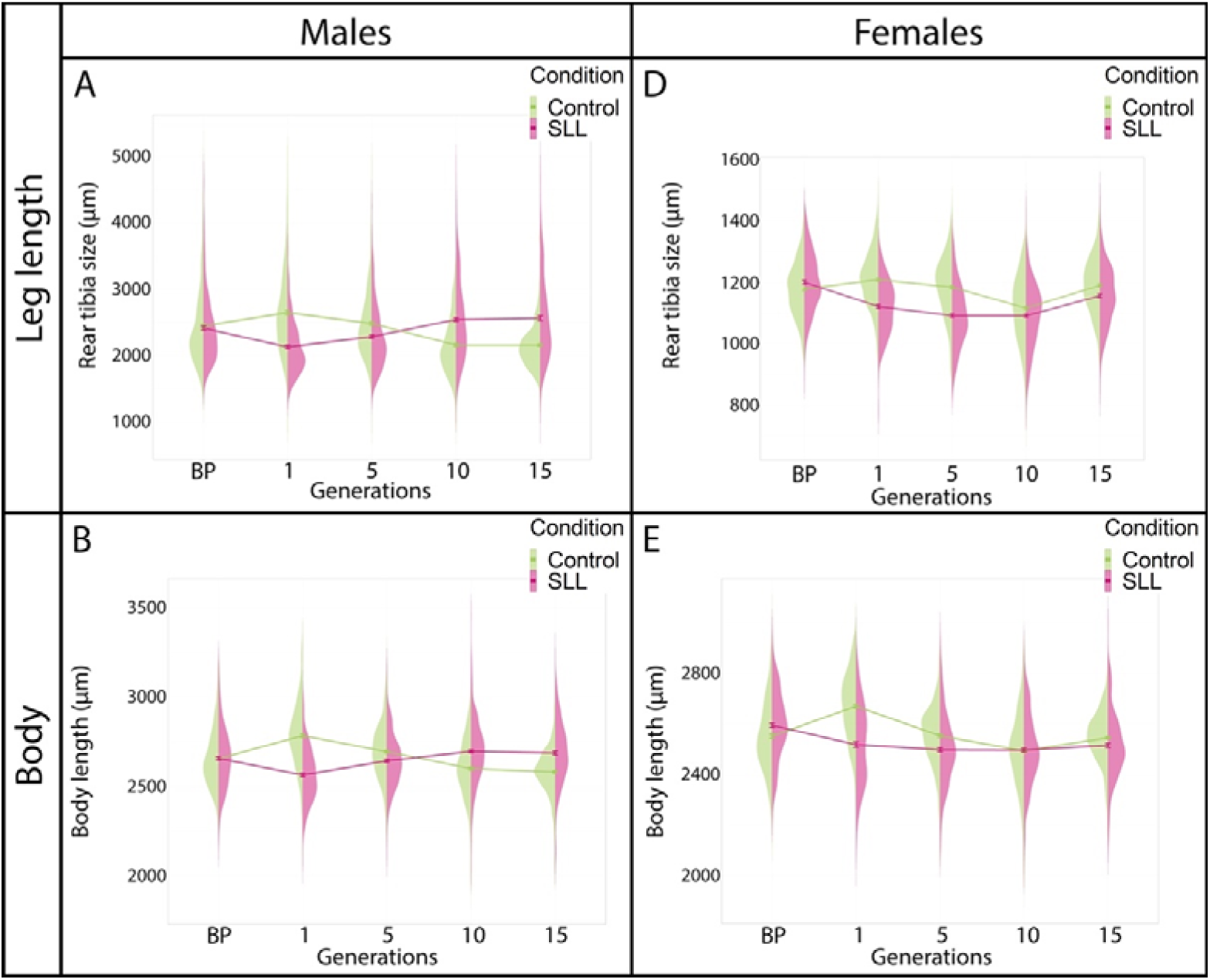
Evolution of rear leg length and body length in males (**A, B**) and females (**C, D**) following male-limited selection in favor of dominance, the mean ± standard-error is represented. BP: base population. Data from the base population (BP), generations 1, 5, 10 and 15 are represented.

Females from the selection line also responded to the standard, less rich, diet since they had legs 6.8% (p < 0.0001) and bodies 2.5% smaller (p = 0.0001) than females from the base population (**Figure 2C-D**). Female rear leg length and body length responded differently to the experimental conditions than males. In both lines, mean rear leg length decreased till generation 10 (p = 0.1809), and then increased from generation 10 to 15 (p = 0.1440) to reach a similar length compared to the base population (**Figure 2D**). In the selection line, the variation in females’ leg length was not impacted by the experimental condition over generations (all p-values > 0.300). Contrastingly, there was an increase of variation at G5 and G10 up to 33% (30-36%, p < 0.0001), then it reduced between G10 and G15 to reach the same value of dispersion as the base population (p = 1.00). Between the two lines, the evolution of mean body length follows the same pattern as the rear legs (**Figure 2**), they diverged until generation 10 (p = 0.4424) and stayed similar at generation 15 (p = 0.1431). However, females of both lines were smaller than the base population. Interestingly, variation of body length decreased by 40% (p < 0.0001) between the base population and generation 15 in the control line, whereas in the selection line body length variation was similar to the base population (**Figure 2**). This result points to a possible genetic correlation in body size between the sexes.

### Selection favouring dominant males increases mean, variance and allometric coefficient of weapon

To disentangle the effects of sexual selection from nutritional environment, samples of individuals from both lines at generation 15 were reared under identical dietary conditions (standard or rich) during nymphal development. Within each line, males raised on a rich diet had longer legs and body than males raised on standard less rich diet (Compare **A** and **B** in **Figure 3; Supplementary Figure S1**). Interestingly, the effect size of diet was greater in the selection line, in which males under the rich diet had rear legs 20% longer (p < 0.0001), whereas in the control line they only differed by 5.5% (p < 0.0001) (**Figure 3A-B**). This further supports the conclusion that phenotypic plasticity will expand standing genetic variation into higher phenotypic variation.

**Figure 3.**
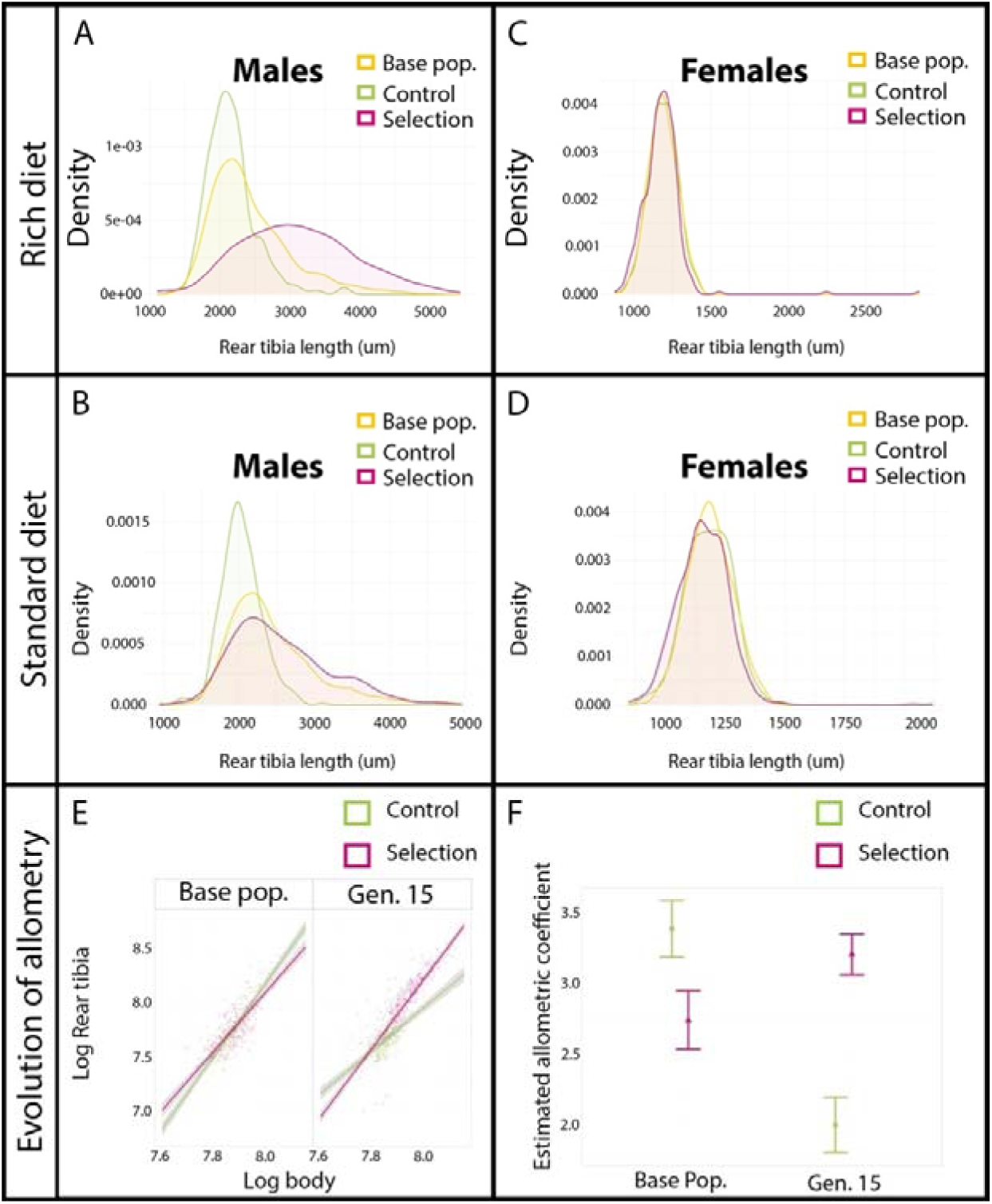
Comparison of mean, variation (**A-D**) and scaling relationships between lines at generation 15 to the base populations subjected to the same nutritional treatment. (**A-B, E**) data for males and (**C-D, F**) data for females. Green represents the control lines, purple the selection lines and yellow the base population. Note that all replicates of the base population, representing both control and selection base populations, are combined. Analyses with base populations are kept separate can be found in **Supplementary Figure 1**. (**E-F**) Scaling relationships of log-transformed data between rear legs and body sizes estimated in males (**E**) fed on rich diet from the two lines from the base population and generation 15. (**F**) Allometric coefficient of log-transformed data between rear leg and body lengths estimated in males fed on rich diet from the two lines from the base population and generation 15, bars represent 95% confidence intervals. Control lines (green and circle), selection lines (purple and triangle).

When we compared rear leg and body length across generations and treatments, we found that lines from generation 15 fed on rich diet had legs 27.9% longer (p < 0.0001) and bodies 5.8% longer than males from the base population (**Figure 3A; Supplementary Figure S1**). Conversely, males from control lines have rear legs 11.5% shorter and bodies 2.8% shorter than the base population (**Figure 3A**). Males from selection lines had rear legs and bodies 41.2% (p < 0.0001) and 8.9% (p < 0.0001) longer than control lines at generation 15 (**Figure 3A**). Moreover, the rear legs of males from the selection line were 43% (p < 0.0001) more variable than males from the control line at generation 15 (T**able 1**). The differences in mean and variation held but were less pronounced in the standard less rich diet (**Figure 3B**). These results indicate that, under selection favouring male dominance, both the mean and variance of male weapon increase, whereas in its absence, they decrease.

**Table 1.** Coefficients of variance (CV) of rear leg and body length in the males of various populations and treatments in *M. longipes*. Note that adults from the two natural populations were captured in the wild and their nutritional status is unknown. The rich and poor diet that the lines were subjected to during nymphal development was described in Toubiana and Khila 2019.

|  | Rich diet |  | Poor diet |  | Natural pop. |  |
| --- | --- | --- | --- | --- | --- | --- |
|  | CV Rear legs | CV Body | CV Rear legs | CV Body | CV Rear legs | CV Body |
| Brazil natural pop | na | na | na | na | 29.2 | 8.8 |
| French Guiana natural pop | na | na | na | na | 24.6 | 7.8 |
| Line LH | 16.8 | 4.7 | 18.4 | 4.5 | na | na |
| Line LL | 12.8 | 3.7 | 16.2 | 5.8 | na | na |
| Line SH | 16.5 | 5.1 | 11.9 | 5.1 | na | na |

Under the same nutritional treatments at generation 15, there was no difference in the length of female rear legs or body between the lines in both diets (all p-values > 0.08; **Figure 3C-D**). However, within each experimental line, females fed on a rich diet showed a modest increase in rear leg length (1.1% longer in rich compared to standard in the control lines (p = 0.0296) and in the selection line 1.4% longer (p = 0.0178). Altogether, these results indicate that directional sexual selection not only fails to reduce variation in male weapon size, but it is rather necessary to maintain it, and that this effect is specific to males in *Microvelia longipes*.

The allometric coefficient of covariation between male rear legs and body also responded to selection favouring dominant males. By generation 15, the allometric coefficient of the control line had decreased relative to the base population (**Figure 3E-F**); (2.0 for the rich diet, p < 0.0001) (2.61 for the standard diet and 2.81 for the rich diet, p < 0.0001). In contrast, males from the selection line maintained a high allometric coefficient (**Figure 3E-F**). Under the standard diet, their slope did not differ significantly from that of the base population (slope: 3.50, p = 0.071), whereas under the rich diet the slope was higher (slope: 3.54, p = 0.011). At generation 15, the allometric coefficient was also significantly higher in the selection line compared to the control line in both diets (p < 0.0001). In females, we found no difference in allometric coefficient between the lines and over generations (**Supplementary Figure S2 and S3**) except the control line fed on rich diet at generation 15 that had a slightly higher slope than females from the base population (slope: 1.85 p < 0.0001). These data indicate that the two treatments diverged in opposite directions from the ancestral population and that selection favouring dominant males is necessary to maintain a high mean and variance of weapon size (**Figure 2**; **Figure 3**).

### Selection favouring male weapon imposes fecundity costs on females

Sexually antagonistic selection is a powerful evolutionary force that can act in opposite directions in males and females when traits are genetically correlated between the sexes (Arnqvist and Rowe, 2005). We therefore hypothesized that the maintenance of high variation in male weapon, in the face of directional sexual selection, could be the consequence of costs to females. To test this hypothesis, we compared female fecundity, a primary fitness component, between selection (long legs/high variance) and control lines (short legs/low variance). Females from selection lines laid 23% fewer eggs than females from control lines, regardless of the origin of the males that they mated with (males from selection or control lines) (**Figure 4A**). The month of the experiment accounted for a moderate proportion of variance in egg number (ICC = 0.39), indicating that while it contributes to variability, most variation occurs among replicates within a month.

**Figure 4.**
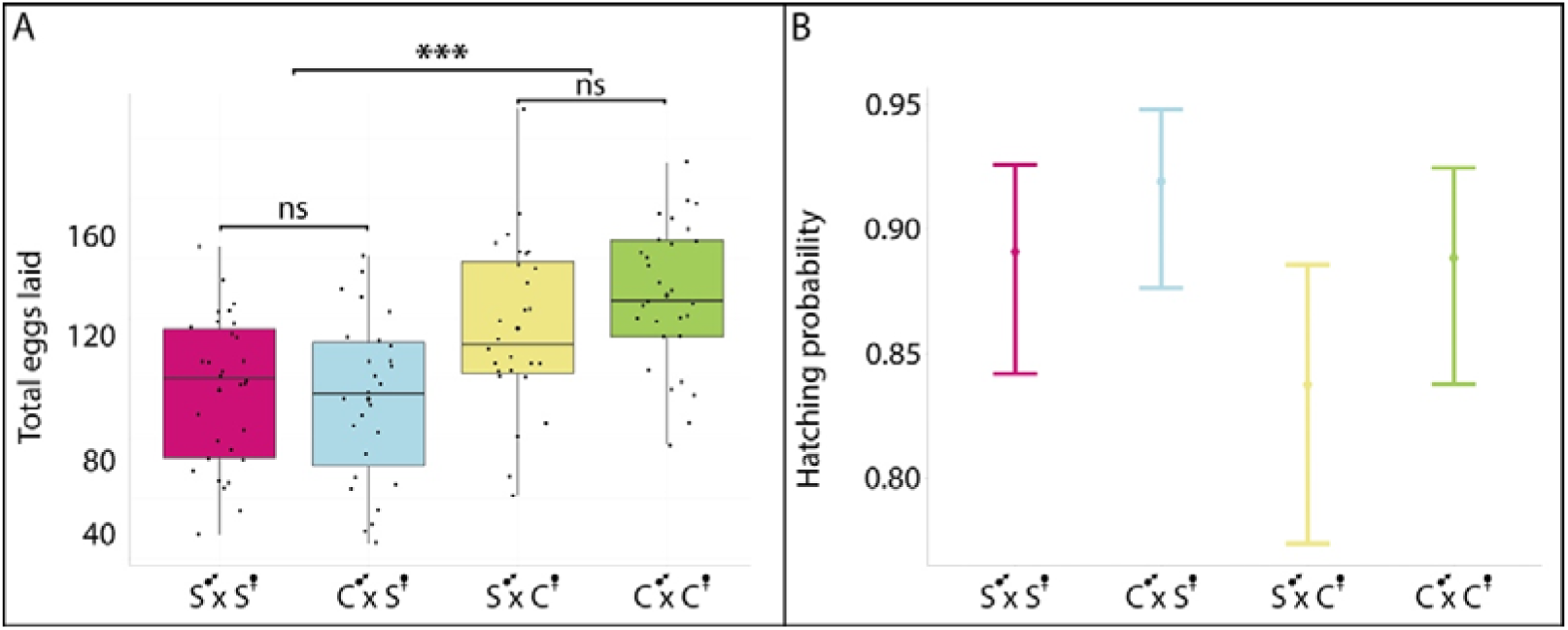
Estimation of fecundity in females through egg production and rate of egg hatching. (**A**) Raw number of eggs laid by females from crosses with males randomly chosen from control and selection lines. (**B**) Probability of egg hatching for the four crossings. Bars represent 95% confidence intervals for model estimated probability.

Probability of egg hatching did not differ between treatments (**Figure 4B**), indicating that the fecundity costs experienced by females from selection lines resulted from reduced egg production rather than reduced egg viability. In addition, the month of the experiment explained only about 22% of the total variance in hatching success (ICC = 0.22), with the majority of variability occurring among replicates within a month. These results demonstrate that sexual selection favouring dominant males generates fecundity costs in females. Therefore, sexually antagonistic selection acts in opposition to directional sexual selection maintaining both large and small males in the population.

## Discussion

We have shown that phenotypic plasticity can expand dramatically the variation in male weapon, and that this effect requires the presence of high standing genetic variation. We have also shown that selection favouring dominant males does not erode, but rather expands variation of male weapon size. This male limited selection generated costs to females in the form of reduced fecundity, highlighting the joint role of directional selection and sexual conflict in maintaining variation. These findings provide a compelling set of advances to our understanding of how variation in exaggerated secondary sexual traits is maintained in natural populations.

### Role of phenotypic plasticity and genetic variation in trait exaggeration

Our reaction norm experiments reveal that phenotypic plasticity alone is insufficient to explain the extreme variation observed in male rear leg length in *Microvelia longipes*. While nutritional variation contributes to expanding phenotypic variation, the dramatic reduction in variation observed in inbred lines and reaction norm experiments highlights the importance of standing genetic variation for plasticity to be able to inflate phenotypic variation in this male weapon. These data align with the genic capture model (Rowe and Houle, 1996), which posits that condition-dependent traits capture genetic variance underlying condition, thereby maintaining high phenotypic variation. However, our results extend this framework by demonstrating that directional sexual selection is not only compatible with the persistence of variation but is necessary for its maintenance in this system. This challenges the conventional view that strong directional selection should erode variation (Falconer and Mackay; Kaufmann et al., 2023; Lande, 1980; Lynch and Walsh, 1998), instead suggesting that selection, in particular for complex traits, may generate a broad range of allelic combinations in the population and sustain variation in the trait.

The experimental evolution lines provided a unique opportunity to disentangle the effects of selection and plasticity. In the context where directional sexual selection is absent or reduced (control lines), variation in male leg length collapsed regardless of nutritional treatment. Conversely, lines subjected to directional selection for exaggerated leg length not only maintained but increased variation in this trait compared to the starting population and to the control lines. This result suggests that directional sexual selection may be part of a mechanism that actively contributes to the accumulation of genetic polymorphism in condition-dependent traits, possibly by favouring genetic combinations that confer advantages to males through dominance of egg-laying sites and access to females. The increased allometric coefficient in selection lines further supports this conclusion, indicating that directional sexual selection amplifies the scaling relationship between weapon size and body size, reinforcing the size of the weapon, more than body size, as a primary target of selection.

### Sexually antagonistic selection as a hidden cost of male dominance

A key insight from our study is the fecundity cost imposed on females in lines subjected to male-limited selection for exaggerated leg length. Females from selection lines laid 23% fewer eggs than those from control lines, regardless of the male’s origin. This demonstrates that sexual selection on males can have negative effects on female fitness, a hallmark of sexually antagonistic selection. Such conflicts arise when traits that enhance male reproductive success incur costs for females, often due to shared genetic architecture between the sexes (Rice, 1984). How this correlation operates at the genomic level remains to be tested. Developmental pathways involved in growth and patterning are known to be highly pleiotropic and often expressed across a number of tissues and developmental stages (Jee et al., 2026; Mackay and Anholt, 2024; Mauro and Ghalambor, 2020). Pleiotropy, the phenomenon whereby a single genetic variant influences multiple biological processes, is now recognized as a pervasive feature of genetic architecture (Mackay and Anholt, 2024; Paaby and Rockman, 2013). In *Microvelia longipes*, pleiotropy may underlie the maintenance of variation in male leg length by linking this trait to other fitness-related pathways, such as those governing body size or fecundity. This is likely as, in this species, the gene BMP11 has been shown to regulate leg length, body size and aggressive behaviour in males as well as body size in females (Toubiana, 2021). Furthermore, a number of studies have shown that pleiotropic genes are often enriched in fundamental cellular pathways (e.g., cell proliferation, DNA repair, and signaling) (Shikov et al., 2020), which could explain how selection on male weapon imposes costs on female fecundity through shared genetic pathways.

In this context, our results support the hypothesis that sexually antagonistic selection acts as a balancing force, counteracting directional selection for trait exaggeration in males. Many of the alleles increasing leg length in males may also decrease egg production in females. Therefore, by reducing female fecundity, selection on male weapon may limit the spread of alleles favouring extreme leg length, thereby maintaining genetic variation in the population. This mechanism complements the genic capture model, as it provides a selective counterbalance that prevents the fixation of extreme traits. The lack of difference in egg hatching success between treatments further suggests that the cost is primarily mediated through reduced egg production, rather than viability, aligning with previous studies on sexually antagonistic fitness effects (Chippindale et al., 2001).

### Phenotypic plasticity as an amplifier of genetic variation in a tug-of-war framework

A central insight from our study is that phenotypic plasticity does not merely respond to environmental cues but actively inflates the genetic variation underlying exaggerated secondary sexual traits, particularly when these traits are subject to opposing selective pressures. Theory predicts that adaptive phenotypic plasticity evolves to reduce evolutionary conflict between the sexes, with the sex avoiding costs being the one that does not express plasticity (Day and McLeod, 2018). Our results in *Microvelia longipes* align with this prediction. Male rear leg length is simultaneously pulled in two directions: directional sexual selection favours longer legs for male dominance, while sexually antagonistic selection imposes fecundity costs on females, creating a genetic tug-of-war. Females may incur reduced fitness costs if males, through plastic responses to nutrition, express high trait values while carrying female-beneficial alleles. As such, females in selection lines, which carry the genetic load of alleles favouring longer legs in males, experience reduced fecundity. This cost creates balancing selection pressure, preventing the fixation of extreme alleles, as males with alleles for longer legs (higher dominance) may produce females with lower fecundity and thus lower fitness. Phenotypic plasticity allows males with intermediate or shorter legs to achieve higher trait exaggeration in response to environmental conditions (e.g., diet), achieving some competitive advantage without transferring alleles of lower fecundity to their daughters. This phenomenon, where selection on male weapon reduced female fecundity, has also been observed in the bulb mites and in beetles (Harano et al., 2010; Plesnar Bielak et al., 2014), suggesting that this mechanism may be more general than previously thought. This plastic compensation ensures that genetic variation is not only preserved but also expressed across a broader phenotypic spectrum. Therefore, phenotypic plasticity evolves as a mechanism that reduces conflict while simultaneously acting as a catalyst in this dynamic, translating the resulting genetic variation into broad variation in male weapon size (**Figure 5**).

**Figure 5.**
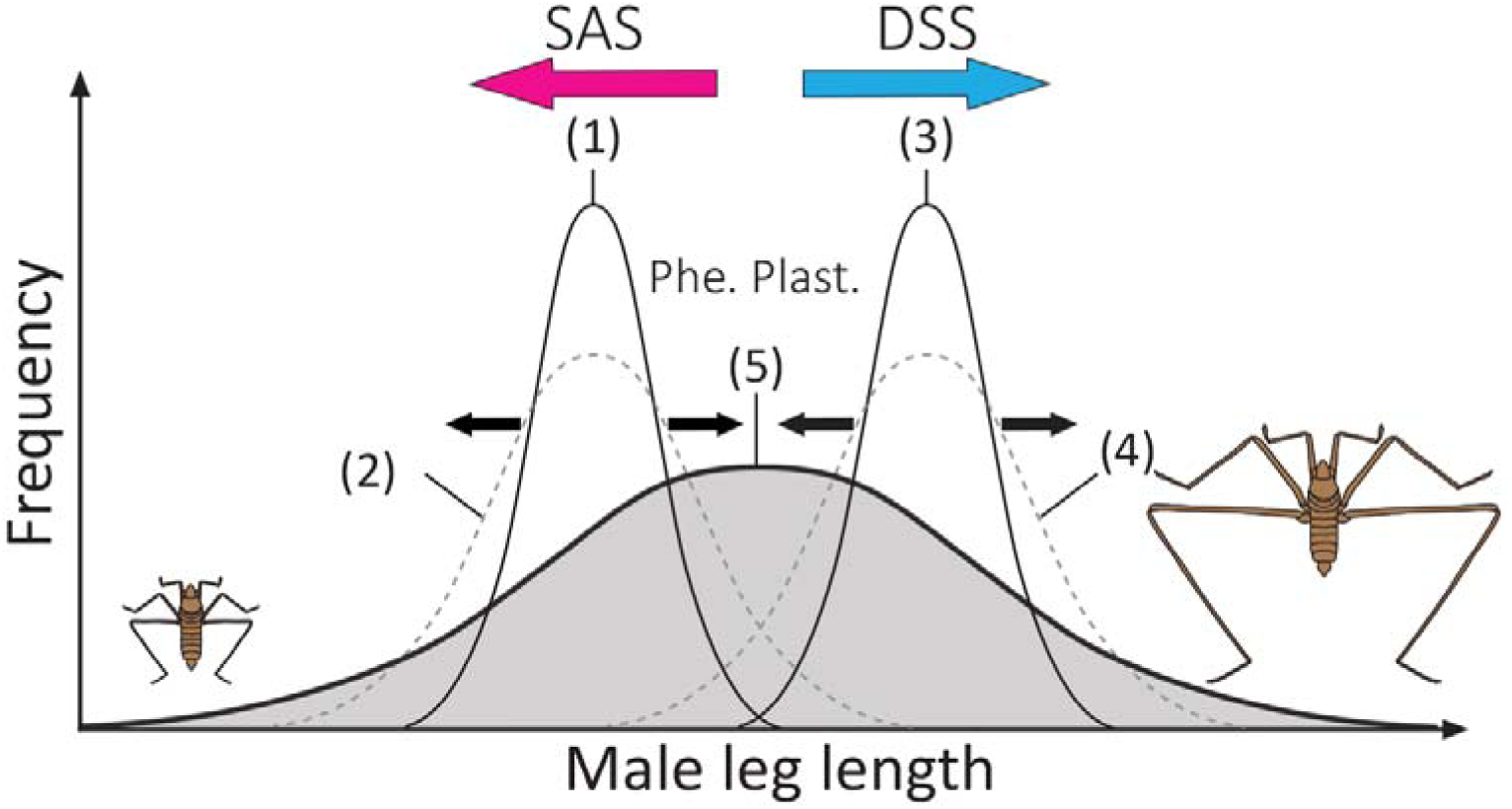
A mechanism underlying the evolution of extreme variation in exaggerated secondary sexual traits. (**1**) In the absence of directional sexual selection and sexual conflict, exaggerated traits are not expected to evolve and phenotypic variation remains narrow. (**2**) Under these conditions, phenotypic plasticity alone (black arrows in **1**) cannot generate either trait exaggeration or extreme variation. (**3**) Directional sexual selection alone can drive the evolution of exaggerated trait values (blue arrow), but phenotypic variation is expected to remain limited. (**4**) In this scenario, phenotypic plasticity may increase variation slightly (black arrows in 3) but cannot account for the extreme variation observed in many sexually selected traits. (**5**) The interaction between directional sexual selection (blue arrow), sexual conflict (red arrow), and phenotypic plasticity (black arrows) generates the extreme phenotypic variation characteristic of exaggerated sexually selected traits.

Our experimental results demonstrate that plasticity expands the range of phenotypic expression for a given genotype, allowing genetically diverse males to achieve even greater variation in leg length under favourable conditions (e.g., rich diet). For instance, males in the selection lines raised on a rich diet exhibited a 20% increase in leg length compared to those on a standard diet, whereas the effect was only 5.5% in control lines. This suggests that plasticity magnifies the phenotypic consequences of genetic differences, particularly when selection is strong. In this way, plasticity does not just mask or reveal genetic variation, but rather amplifies it providing a mechanism for the persistence of individuals with extreme and intermediate trait expression despite opposing selective forces.

Finally, the allometric scaling of leg length relative to body size in selection lines, where the slope of the relationship increased under rich diet, further illustrates how plasticity can expand the phenotypic landscape. By allowing genotypes to express a wider range of trait values under different conditions, plasticity effectively inflates the variance generated through the opposing effects of directional sexual selection (favouring males with long legs) and sexually antagonistic selection (favouring more fecund females). This may explain why, despite strong directional selection for longer legs, variation in *M. longipes* remains so large, as plasticity converts this genetic variation into large variation in male weapon, thus providing material for both selection and conflict to operate.

### Sexual selection and the maintenance of variation

The persistence of variation in exaggerated secondary sexual traits has long been a paradox in evolutionary biology. Our study provides empirical evidence that directional sexual selection is required to maintain, rather than deplete, genetic variation in such traits. This finding contrasts with theoretical predictions that strong directional selection should reduce variation. However, our results align with emerging evidence that condition-dependent traits may capture and amplify the underlying genetic variation (Parrett et al., 2022), particularly when selection targets multiple loci contributing to condition.

The divergence between the selection lines and the control lines in both mean and variance of leg length suggests that selection favours not only the exaggeration of the trait but also the diversification of male strategies. In *M. longipes*, long-legged males dominate reproductive sites, but the persistence of smaller males, despite their lower dominance, implies that alternative strategies (e.g., sneaking or satellite behaviours) may be viable. This has been confirmed through observation in *M. longipes* where small males adopt, often successfully, sneaking behaviour and manage to fertilize a significant number of eggs (Toubiana and Khila, 2019). This observation could also constitute an additional mechanism that contributes to the maintenance of high variation of male weapon in this species.

### Implications for the evolution of trait exaggeration

Our findings refine several cornerstones of evolutionary theory. First, they demonstrate that directional sexual selection does not inevitably lead to the erosion of genetic variation in exaggerated traits. Instead, selection with its various facets can maintain high levels of genetic variation, which provides the material for phenotypic plasticity to inflate into the broad phenotypic variation often observed in exaggerated sexually selected traits. Second, our study highlights the importance of sexually antagonistic selection as a mechanism that counter-balances directional sexual selection. By imposing fitness costs on females, sexual selection on male traits can create a dynamic tug-of-war that prevents the fixation of extreme phenotypes, thereby maintaining polymorphism. This aligns with the growing recognition that intralocus sexual conflict (where males and females share genetic variation but have opposing fitness optima) is a widespread and powerful evolutionary force (Arnqvist and Rowe, 2005). Such conflicts may explain why many secondary sexual traits exhibit such remarkable variation, even in the face of strong directional selection.

Finally, our results underscore the need for integrative approaches in evolutionary biology. The interplay between genetic variation, phenotypic plasticity, and selection cannot be fully understood in isolation. Future theoretical and empirical work should aim to incorporate these interactions into models of trait evolution, particularly for systems where sexual selection is strong and traits are highly condition-dependent. This could reveal new insights into the evolution of complexity, the origins of novelty, and the limits of adaptation.

### Conclusion: a mechanism for maintaining extreme variation in male weapon size

Combined, our findings reveal a mechanism underlying the evolution of extreme phenotypic variation (**Figure 5**). Directional sexual selection, driven by male–male competition, promotes the evolution of exaggerated male weapons by favouring alleles that increase trait size. In contrast, sexually antagonistic selection arising from fecundity costs in females prevents the fixation of these same alleles, thereby maintaining genetic variation within the population. Under this scenario, low, intermediate and high levels of trait expression are maintained within the population. Fitness costs incurred by males bearing small traits may be offset through the production of more fertile daughters, whereas the reproductive advantages of males with exaggerated traits may be counterbalanced by the reduced fertility of their daughters. Phenotypic plasticity can subsequently amplify this standing genetic variation and blur the boundaries between intermediate and extreme phenotypes, resulting in the broad distributions of trait values observed in exaggerated secondary sexual traits. Similar patterns have been reported in other systems. For example, in the broad-horned flour beetle and in the bulb mites, selection favouring increased male weapon size resulted in reduced lifetime fecundity in females, whereas selection against weapon exaggeration increased female fecundity (Harano et al., 2010; Okada et al., 2021; Plesnar Bielak et al., 2014). More broadly, our study highlights how the interaction between genetic conflict, environmental sensitivity and directional selection can be a powerful mechanism generating extreme phenotypic variation in exaggerated secondary sexual traits.

## Materials & Methods

### Study organism

*Microvelia longipes* is a gregarious species that lives in small temporary puddles filled with rainwater in tropical regions of South America (Andersen, 1982; Toubiana and Khila, 2019). Females lay their eggs on debris of twigs and dead leaves floating on the surface of water, that we refer to as egg-laying sites (Toubiana and Khila, 2019). Males guard these egg-laying sites, signal to attract females, and fight using their 3^rd^ pair of legs as weapons to fend off competitors (Toubiana and Khila, 2019). Males with long 3^rd^ pair of legs are favoured by intrasexual competition which has led to the evolution of exaggerated leg length (Toubiana et al., 2021; Toubiana and Khila, 2019).

### Experimental evolution

A schematic overview of this experiment is provided in **Supplementary Figure S4**. A wild population composed of 50 females and 41 males captured in 3 locations in French Guiana in April 2023 (GPS coordinates: 4.30275, -52.13956; 4.04089, -52.02277; 3.90165, -51.81666) was allowed to expand for one generation in the laboratory to generate a base population. From this base population, eight replicate experimental evolution populations were established, four subjected to selection favouring dominant males with long legs (Selection line), and four in which sexual selection was relaxed (Control line). Each population was founded by 260 recently emerged adults from the base population (130 random females and 130 random males). Generations were not allowed to overlap.

#### – Selection regime for dominant males

In *M. longipes*, males compete for egg-laying sites and large males almost always win (Toubiana and Khila, 2019). We therefore based our selection regime on this mating system by reducing the number of egg-laying sites to 1 site per 10 males to exacerbate competition that favours dominant males (Emlen, 2008; Toubiana and Khila, 2019). The size of these sites was also reduced so that only one male at a time can defend it (Emlen, 2008) (1 cm x 0.10 cm). In addition to intensified competition, we also sought to increase selection efficiency by limiting the effect of phenotypic plasticity that hides the genotype from selection (Crispo, 2008; Grenier et al., 2016; West-Eberhard, 2003). We therefore subjected the selection line to a standard diet (frozen crickets) during the nymphal stages (nymphal instars 1 to 5). This treatment, in our experience, is less rich (likely due to the fact that frozen crickets rot quicker than freshly euthanised crickets) and is meant to enhance the congruence between the genotype and the phenotype (Crispo, 2008; Grenier et al., 2016; West-Eberhard, 2003). The nymphs from the selection line were therefore fed once a day with frozen crickets, in the same way as species stock are maintained in the laboratory.

#### – Control line

Control replicates were maintained in conditions of low or no competition for egg-laying sites. They were provided with Styrofoam stripes (2 x 8 cm), which are 160 times larger than the egg-laying sites provided to the selection lines. The size of these floaters makes it impossible for males to defend. This ensures that females lay eggs without having to copulate with a dominant male. In addition, the nymphs from control lines were fed *ad libitum*, twice a day with fresh crickets, in order to amplify phenotypic variation.

For both lines, once the nymphs reached adulthood, they were fed with the same rich diet. The populations were maintained at 28-30°C, 50-60% humidity and under an artificial photoperiod of 14 hours of daylight in water tanks (37x54x21 cm). Eggs laid on the egg-laying sites (Selection line) or large Styrofoam floaters (Control line) were allowed to develop to constitute the next generation. The adults were allowed to interact freely until there were enough nymphs to constitute the next generation (∼260 individuals for each population), then they were photographed for subsequent phenotyping and sacrificed in absolute ethanol before being frozen at -70 °C (**Supplementary Figure 5**).

To disentangle the effects of selection regime from developmental nutrition, individuals from both lines were reared under identical nutritional conditions during G15. For each replicate of each line, nymphs were randomly assigned to a diet (*Ad libitum* diet or standard diet).

From generation 15 on, both lines are maintained at a less intensive level: smaller tanks, smaller population, fed with standard diet (once a day with frozen crickets), but we maintained competition by providing small egg-laying sites to the selection lines and large Styrofoam floaters to the controls.

### Phenotyping

Due to a large number of individuals per generation (∼2 000 across all replicates), we chose a group imaging method that is compatible with quick but reliable measurements (**Supplementary Figure S5**). This method is dependable because the insects adopt a stereotypical posture naturally on the water surface (**Supplementary Figure S5**). At each generation, we subdivided each population in multiple groups of approximately 50 living animals, and pictures were taken using a vertically placed Nikon D7200 camera with AF-S Micro-NIKKOR 105 mm lens. Length of the body, the tibia and the femur of both sexes were measured for the wild population captured in 2023 and for all replicates from the base population and at G1, G5, G10 and G15 using the software ImageJ (v1. 53t). Body length was measured from the tip of the head to the tip of the genitalia. Tibia length from the joint with the tarsus to the joint with the femur, and femur from the joint with the tibia to the joint between the femur and the side of the body (see landmarks in **Supplementary Figure S5**).

### Fecundity assay

To test if selection favouring dominant males had any impact on female fitness, we tested fecundity of the females in both control and selection lines. Four types of crosses were set up (male SLL line x female SLL line; male Control line x female SLL line; male SLL line x female Control line; male Control line x female Control line), each with thirty biological replicates consisting of a single couple each. For each replicate, one male and one female were allowed to mate in a plastic container (14x5x8 cm) filled with water, and two egg-laying sites were made available (each: 1.5 cm x 0.1 cm). Egg-laying sites were changed every two days during 12 days, and the total number of eggs laid and the number of eggs that hatched were counted 7 days after the first hatching, (embryonic development takes about 5 days). A period of 12 days was chosen, prior to data analysis, as this was the longest period during which no females had died. The average lifespan of an adult female (since emergence) is around 34 days (unpublished data). Individuals were isolated at the 5^th^ instar, to standardise the age of adults and capture most of female lifetime fecundity. When a male died during the experiment, it was replaced by a male from the same line. This experiment was performed twice, once in April 2025 at G23 for the SLL line and G27 for the Control line with 10 replicates per crossing, then in July 2025 at G27 for the SLL line and G31 for the Control line, with 20 replicates per crossing. Egg-laying sites used for the selection lines were smaller, which increased a bit the time required to produce the next generation, thus causing a delay between the two lines.

### Statistical analyses

Statistical analyses were conducted using R software (v.4.2.2) and data were visualized using ggplot2. Log-transformed data were used for scaling relationship comparison, and we performed a linear mixed effects model, the interaction between the generation and the condition as the predictor and the replicates and the person measuring as random effects. We run a similar model for females.

Comparisons for trait distributions were also performed on log-transformed data. We tested if the length of the tibia of the rear leg and the dispersion (proxy of the total leg length (Toubiana and Khila, 2019) changed across generations due to the experimental condition using a generalized linear mixed model with tibia length as the dependent variable, the interaction between the generation and the condition as the predictor and the replicates and the person measuring as random effects (‘glmmTMB’ package in R). Similar models were run for the length of the tibia of the rear leg in females and body length in both sexes. At generation 15, we reared individuals from both lines under identical nutritional conditions to disentangle the effects of selection regime from nutritional treatment. In order to test for the effects of these parameters on tibia and body length on individuals at generation 15, we used a linear mixed-effects model with the length of the body or tibia as the dependent variable, the interaction between the diet and selection regime as the predictor and the replicates and the person measuring as random effects.

The effect of the crosses on the total number of eggs laid by females was tested through using negative binomial generalized linear mixed model with a log link function (“glmmTMB” package in R). The probability of hatching was also compared between crosses using a beta-binomial mixed model (“glmmTMB” package in R). The month of the experiment was included as a random effect for both models. Model adequacy was assessed using DHARMa residual diagnostics, which indicated no remaining overdispersion or deviation from model assumption.

## Acknowledgement

The authors acknowledge support from the PSMN (Pôle Scientifique de Modélisation Numérique) of the ENS de Lyon and from GENCI/TGCC (grant A0110807662) for the computing resources. This work was supported by a MITI fellowship, FRM équipe grant EQU202103012573 and VR 146700220 grant from Swedish research council grant to AK.

