## Supplementary online information for "Sexual conflict, directional sexual selection and phenotypic plasticity jointly drive the evolution of extreme phenotypic variation"

**Author affiliation :**

### Supplementary figures

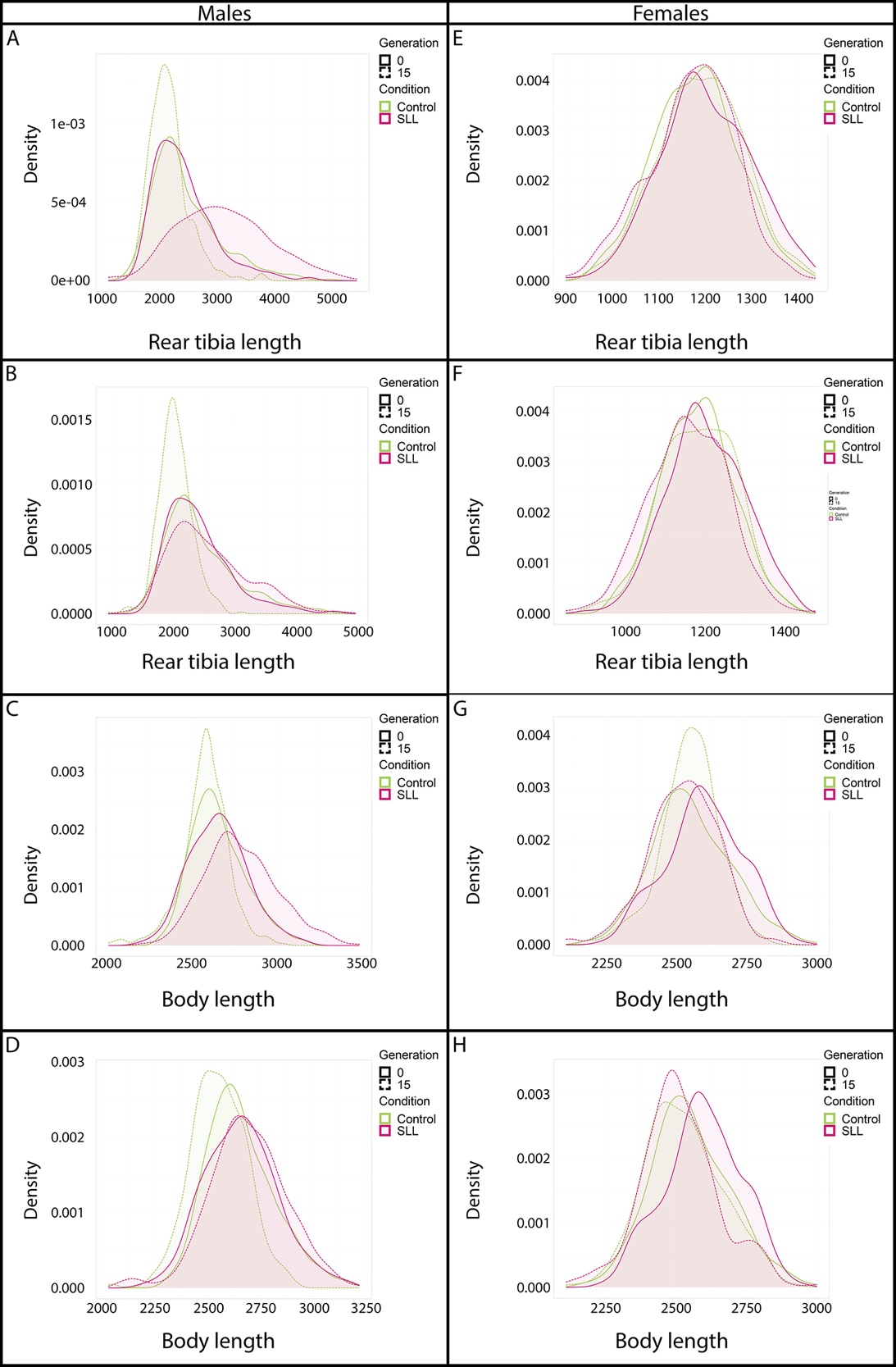

**Supplementary Figure S1.** (A) evolution of the length of the rear leg in females, the mean ± standard-error is represented (B) evolution of body length in females, the mean ± standard-error is represented (C) and (D) distributions of the length of rear leg and body at G15 in *ad libitum* diet, (E) and (F) distributions of the length of the leg and body at G15 standard diet. Green represents control lines, and purple selection lines. Plain lines represent the base populations and the dashed lines generation 15.

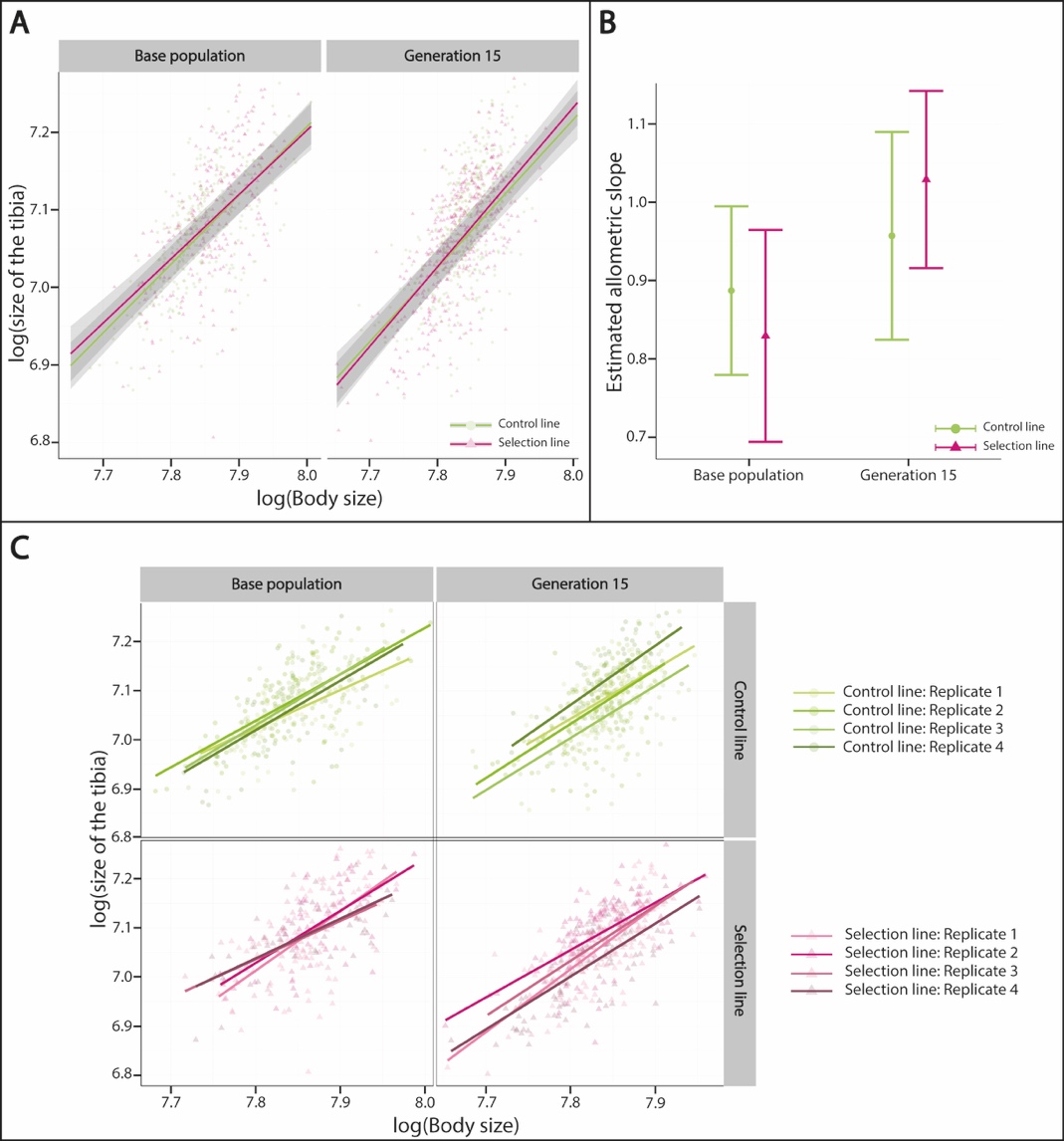

**Supplementary Figure S2.** (**A**) Scaling relationships of log-transformed data between rear legs and body sizes estimated in females fed on rich diet from the two lines from the base population and generation 15. (**B**) Allometric coefficients of log-transformed data between rear leg and body lengths estimated in females fed on rich diet from the two lines from the base population and generation 15, bars represent 95% confidence intervals. Control lines (green and circle), selection lines (purple and triangle). (**C**) Scaling relationships of log-transformed data between rear legs and body sizes estimated in females fed on rich diet from the two lines from the base population and generation 15 for each replicate.

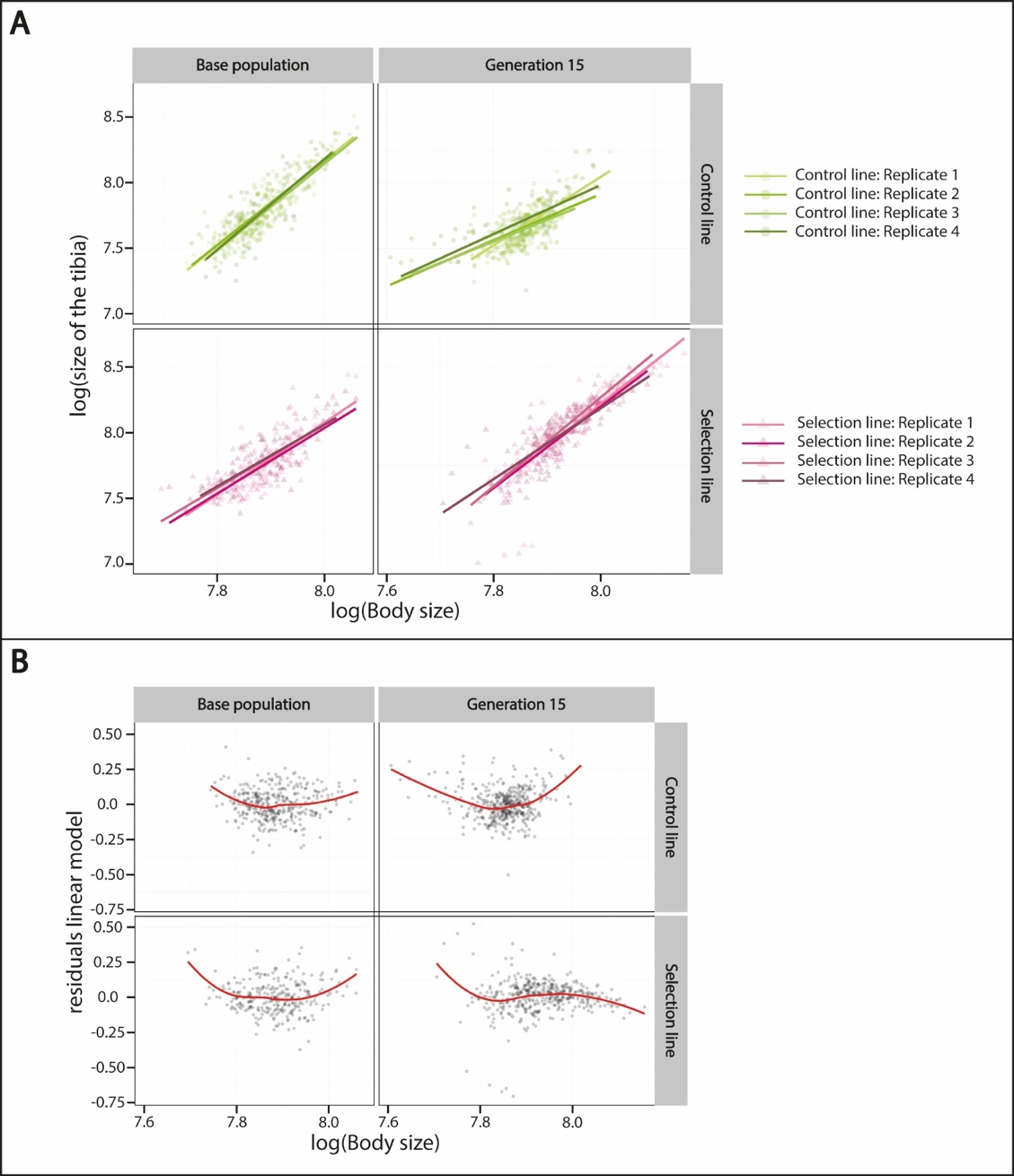

**Supplementary Figure S3.** (**A**) Scaling relationships of log-transformed data between rear legs and body sizes estimated in males fed on rich diet from the two lines from the base population and generation 15 for each replicate. (**B**) Residuals of the linear mixed effects model for males.

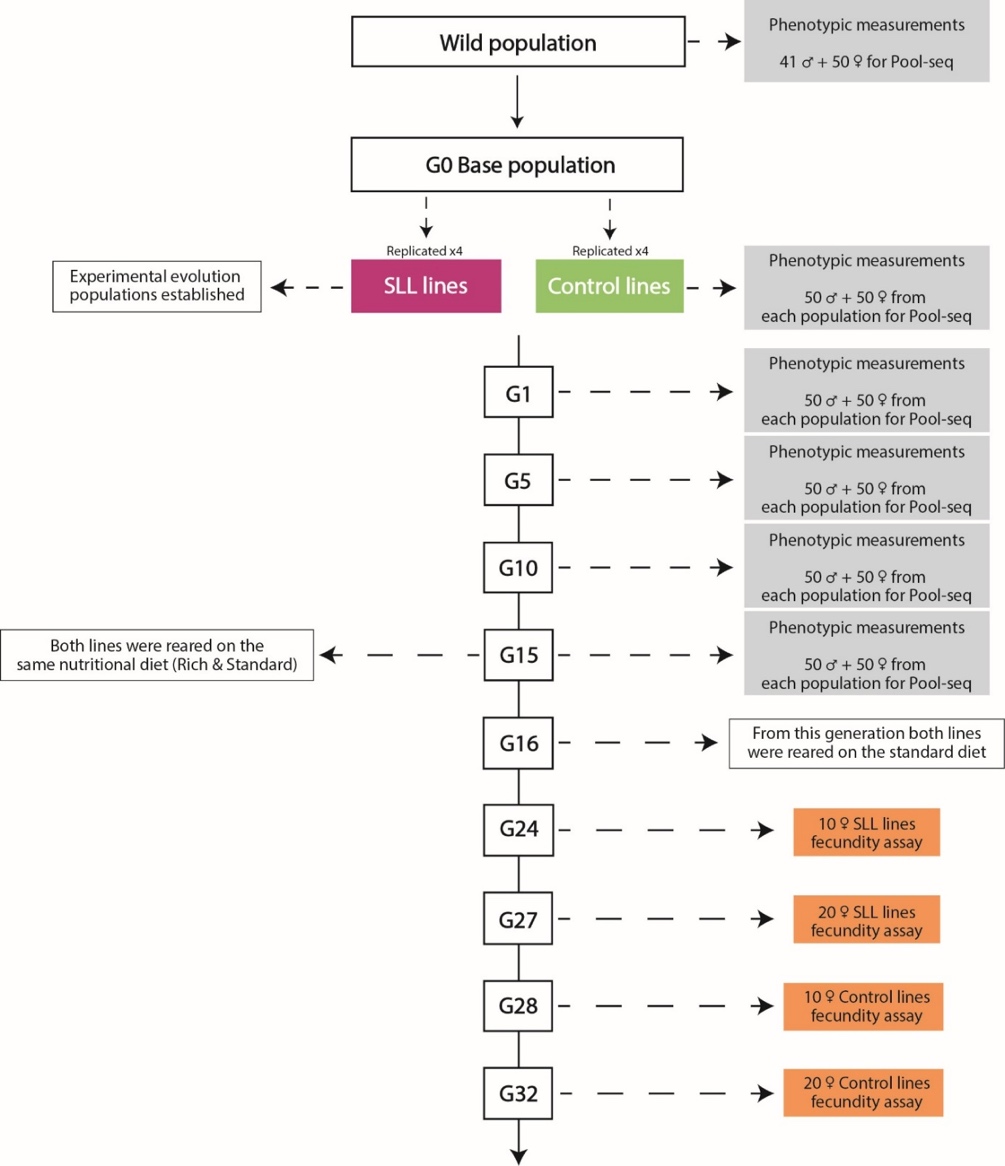

**Supplementary Figure S4.** Schematic overview of the experimental design. Shown are the generations when sampling for resequencing and phenotypic assays were performed (grey boxes). Also shown are the generations at which fecundity assays were performed (orange boxes).

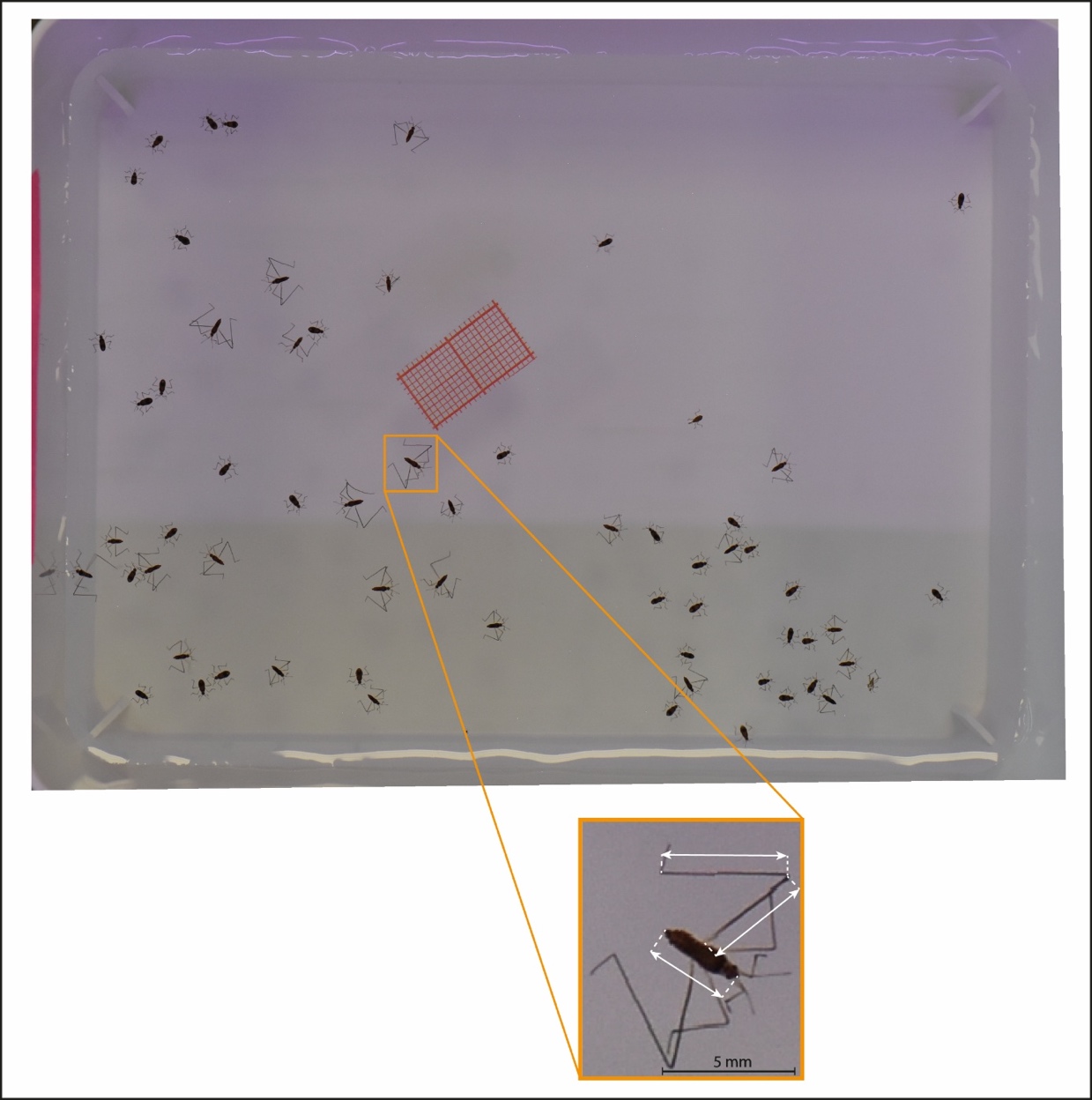

**Supplementary Figure S5.** Representative image of the phenotyping procedure employed to measure the experimental evolution samples. The orange box represents the scale at which the individuals were measured, and the arrows represent the landmarks used to measure body size, tibia and femur length.
